# Chimerism and altruism

**DOI:** 10.1101/2024.07.23.604446

**Authors:** Thomas J. Hitchcock, Manus M. Patten

## Abstract

Chimerism spans the tree of life, from mammals and corals to plants and fungi. In such organisms, individuals contain within them cells and genomes from another once distinct member of the population. This chimeric genetic composition may subsequently alter patterns of relatedness not only between those individuals, but also within them. Consequently, we may expect unique patterns of social behaviour in such species. To explore the social evolutionary consequences of chimerism, here we develop a kin-selection model of a structured population. First, we show how somatic and germline chimerism influence patterns of relatedness and play an important role in modulating social behaviour. Specifically, we find that increased heterogeneity of the soma relative to the germline boosts the opportunity for altruism between individuals. We then explore how differences in chimerism levels within the body may generate within-organism differences in the valuation of social partners and thus foment internal conflicts between tissues and organs. Finally, we show how differences in the development of male and female germlines in chimeras provides a novel source of relatedness asymmetry between maternal-origin and paternal-origin genes. Overall, we find that chimerism introduces additional opportunities for internal conflicts over the development of behavioural phenotypes, most of which have been unexplored by empiricists.

## Introduction

Many individuals carry genetic variation within each of their cells, inheriting potentially distinct alleles from their mothers and fathers. However, some organisms display not only genetic heterogeneity within their cells but also between them, with genetically distinct cellular lineages coexisting in the same body. Most commonly this may arise from somatic mutations that occur through development, known as mosaicism (Otto & Orive, 1995; Reusch *et al*., 2021; Frank, 2022). However, there are also more radical forms, where cells from once distinct zygotes now share the same body, a process known as chimerism (Rinkevich, 2011). Whilst such chimeras are relatively rare, they are found across the tree of life, from corals and monkeys to algae, ants, and even ourselves (Ross *et al*., 2007; Boddy *et al*., 2015; Santelices *et al*., 2017; Chang *et al*., 2018; Guerrini *et al*., 2021; Darras *et al*., 2023). Moreover, new genomic tools have opened new windows into these phenomena across a range of different species, giving us a richer picture not only of the frequency of chimerism, but also its genetic structure within organisms (del Rosario et al. 2024).

Chimerism changes patterns of relatedness and predictions from kin selection in two ways. First, if an individual contains multiple, genetically distinct cell lineages, then this decreased relatedness within the individual may foster internal conflict over reproduction and development (Buss, 1982, 1987). Second, that same cell sharing may simultaneously increase an individual’s relatedness to social partners, and thus potentially promote cooperative behaviours between individuals. For example, it has been suggested that the cell sharing between twins during pregnancy in Callitrichidae monkeys may reduce sibling rivalry, and potentially explain their high levels of alloparental care (Haig, 1999). But should this chimerism extend to germlines it may decrease the relatedness amongst broodmates, potentially having the opposite effect (Patten, 2021).

Most of the theoretical work on chimerism to date has focused on the former, intra-individual social behaviour, while relatively less attention has been given to the latter, inter-individual interactions. Moreover, the work that has been done has focused on interactions within the family (Haig, 1999, 2014a, 2014b; Boddy *et al*., 2015; Patten, 2021; U beda & Wild, 2023). Thus, it is less clear how these results apply to other forms of chimerism, and in different demographic settings. Such extensions are important, as patterns of mating, dispersal, and reproductive skew are all known to influence patterns of relatedness and social behaviour (Lehmann & Rousset, 2010; Cooper *et al*., 2018), and thus it is unclear how they may intersect with the complexities of chimerism.

To address this, here we develop a kin-selection model for chimeric organisms. We consider the potential for an altruistic behaviour to evolve in a structured population and how this depends on chimerism’s ontogeny. We consider how the type and timing of chimerism may modulate the potential for altruistic behaviours between individuals. Moreover, we consider how conflicts of interest between cell lineages may emerge over such social behaviours, leading to a suite of internal conflicts—between genes, cells, and tissues. Finally, we discuss how these results relate to currently understood chimeras and possible future avenues for experimental and theoretical exploration.

## Methods and Results

### Life cycle

We consider a population composed of an infinite number of patches, in which there occurs the following life cycle, depicted in Figure 1a. (1) First, a large number of offspring are produced by *N* adults on the patch, (2) offspring then aggregate into “patchlets” of *n* individuals, producing chimeric individuals. (3) The chimeric individuals then grow and develop, redistributing those cell lineages across the body. (4) Individuals may then disperse between patches, remaining on their natal patch with probability *h*. (5) There is density dependent regulation, leaving *N* adult individuals on the patch. The life cycle begins again. We explore some further variants of this life cycle, with different timings of dispersal in the Supplementary Material.

**Figure 1:**
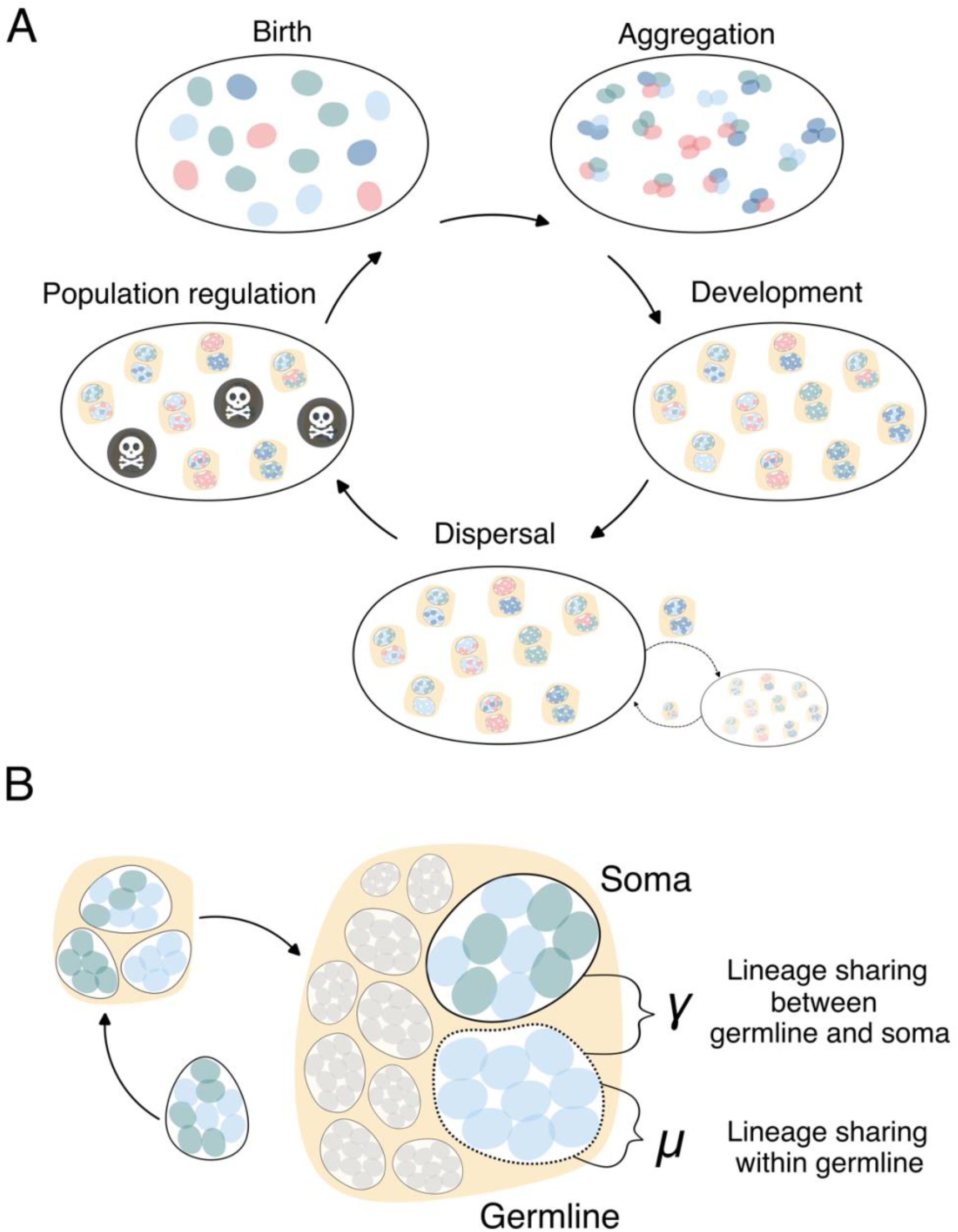
Diagram of chimeric life cycle. (A) The order of key life cycle phases is shown, and (B) development within chimeras is visualised, specifically the parameters *μ*, the probability two cells in the germline descend the same cell lineage, and *γ*,the probability that two cells, one from the germline and one from the soma, descend from the same cell lineage.

**Figure 2:**
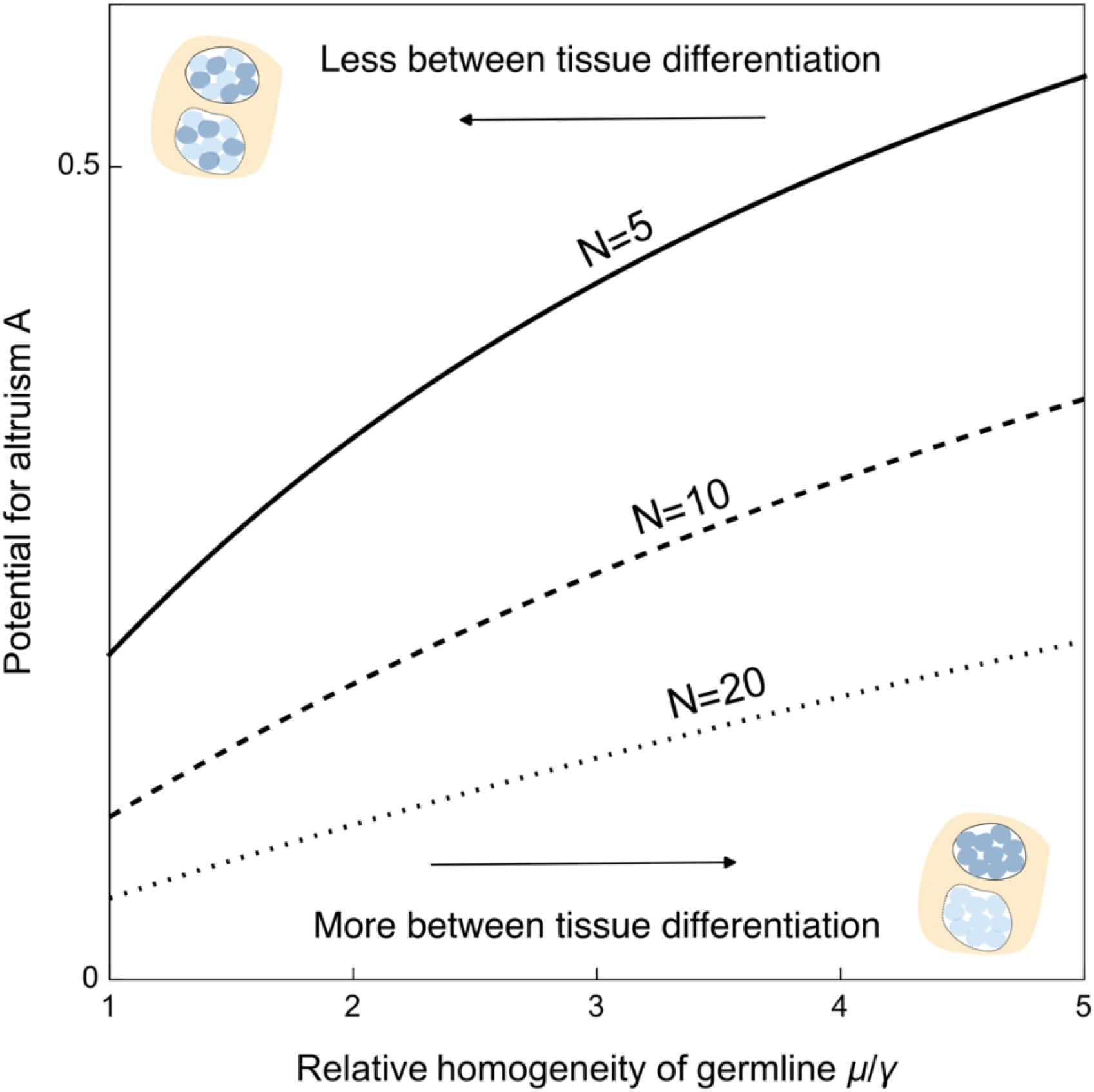
Increasing genetic differentiation between germline and soma promote the evolution of altruism in structured populations. Results for three different patch sizes are shown (*N*), with smaller patches having higher potentials for altruism.

During development, we allow the different cell lineages within our organism to clonally proliferate. We can describe the resulting within-organism genetic structure in the following way. Let the probability that two cells in the germline come from the same cell lineage be *μ*, and let the probability that a cell sampled in the soma and in the germline come from the same cell lineage be *γ* (Figure 1b). This mirrors the approach used to describe clonal proliferation of asexual individuals within patches (Prugnolle *et al*., 2005; Hitchcock, 2024). We can additionally extend this to consider distinct male and female germlines within hermaphrodites by *μ*_ij_ and *γ*_i_, where i and j can be either male or female germlines.

Within this life cycle we consider a social behaviour that modulates the survival of individuals during the juvenile phase *S*(*x,y*). A focal individual’s survival is determined by the phenotype expressed by their soma *x*, and the average phenotype on their patch *y*. We consider this behaviour to be cooperative, such that an increase in the trait value imposes a cost upon the individual’s survival 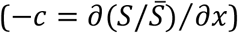 but provides an increase in the probability of survival of individuals on the patch 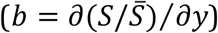. We analyse this model using the direct fitness approach of Taylor and Frank (Taylor, 1996; Taylor & Frank, 1996), assuming that changes in the trait value are small, such that we evaluate at the population average trait of *z*. We then interpret these results in terms of inclusive fitness. Full details can be seen in the Supplementary Material.

### Altruism, relatedness, and the scale of competition

We write the condition for our trait to increase in the form of a Hamilton’s rule, whereby, for the trait to increase in value the direct, marginal cost to self, *C*, must be outweighed by the indirect, marginal benefit to social partners, *B*, where both parts are weighted by the appropriate relatedness coefficients. Assuming that gene copies residing in the somatic tissue control the trait of interest, then the appropriate relatedness coefficients will be between the focal individual’s somatic tissue and the germline of that same individual, *r*_*s*_, and the relatedness between the focal individual’s somatic tissue to the germline of their social partners, *r*_*p*_. This gives us the following condition:

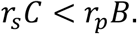

We can further expand this and write the *C* and *B* in terms of marginal effects upon survival, *c* and *b*. As evolutionary success is relative, a change in the fitness of one individual must, ultimately, be compensated by the loss in fitness of another individual. In a structured population, individuals may compete locally, causing some of this fitness compensation to fall upon relatives. Thus, the marginal fitness benefit (or cost) that an individual experiences (−*C,B*) may differ from the marginal survival benefit (or cost) that it received (−*c,b*). We can describe the intensity of this effect with *a*, the scale of local competition, which captures the fraction of that compensation that accrues to local patchmates (who are related by *r*_*p*_) as compared to unrelated members of the population (Frank, 1998; Scott & Wild, 2023). Depending on the life cycle and the timing of regulation and dispersal, *a* may take on different values. Incorporating this, the condition for increase becomes

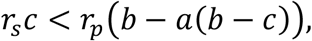

which can then be rearranged into the following form:

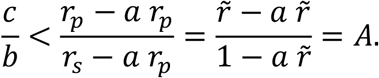

We define the right-hand side as *A*, the potential for altruism (Gardner, 2010). As this quantity increases, the condition for increase becomes easier to satisfy, and thus altruistic behaviours are more likely to invade a population. The potential for altruism can also be seen as a relatedness coefficient which incorporates the relatedness of competitors (Queller, 1994; Scott & Wild, 2023). The inverse of this quantity, *k*=1/*A*, is also used as a measure of how much larger the benefit to the recipient has to be to compensate for the cost to the focal actor (Hamilton, 1963; Patten, 2021).

Inspecting this equation, we can see that increasing the relative relatedness to social partners 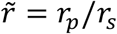, will promote altruism. In the most familiar form of Hamilton’s rule, 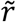 is simply *r*_*p*_, and *r*_*s*_ is taken to be 1. But chimerism may mean that an allele in the soma may not be present in the germline of its own body. By decreasing the relatedness to oneself— or, more accurately, the relatedness of the soma to the germline in a single body— 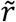 increases, making for a relatively greater return for the soma on the investment in social partners. At the same time, however, chimerism may reduce *r*_*p*_, making it difficult to simply intuit the net effect of chimerism on the evolution of social behaviour.

In addition, reducing the scale of competition, *a*, will also promote altruism. In many classical models, limited dispersal, or viscosity, raises the relatedness to social partners, thus raising 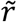, but at the same time heightens the scale of competition, *a*, in such a way that the effect of dispersal neatly cancels and leaves the potential for altruism unaffected (Bulmer, 1986; Taylor, 1992). Whether this cancelation also holds for chimeric organisms has not previously been examined.

### Chimerism and altruism

We now consider how chimerism will shape the relatedness to social partners, and subsequently the potential for altruistic behaviours. Focusing on where there is one phase of dispersal, after our social behaviour but before density regulation (Gardner, 2010), then the relatedness coefficients are:

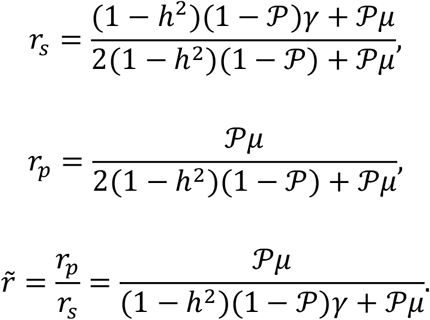

where 𝒫 is the probability that two individuals share the same parent and the scale of competition in this demography is *a*=*h*^2^. Substituting these values into our expression from earlier, the potential for altruism becomes:

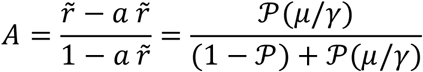

where *μ* is the probability that two cells in the germline come from the same cell lineage and *γ* is the probability a cell sampled in the soma and the germline come from the same cell lineage.

If *μ* and *γ* are both simply 1, as they would be in a non-chimeric organism, then the result simplifies down to 𝒫. If there are *N* breeders with Poisson distributed number of offspring then this would be *A*=1/*N*, which is the classic result initially found by Taylor (1992). Moreover, this result holds for any case where the consanguinity within the germline, and between the soma and germline, are the same (*μ*=*γ*). In such cases, the absolute amount of chimerism would be expected to have no impact upon patterns of interindividual social behaviour. Additionally, across these cases, regardless of the extent of chimerism, the potential for altruism remains invariant to dispersal.

However, *μ* – the similarity within the germline - and *γ* – the similarity between germline and soma - may not be the same. Increasing the ratio of *μ*/*γ* promotes altruism, whilst lowering *μ*/*γ* inhibits altruism. Thus, chimerism has the largest altruism-promoting effect when the germline is relatively homogenous but the soma and germline are genetically distinct. Such a situation may occur if an initially genetically heterogeneous chimera is partitioned through small bottlenecks into germlines and somatic tissues or if an individual’s germline is sequestered pre-fusion and then protected from invasion by the newly fusing cell lineage.

The reason that a greater *μ*/*γ* ratio raises the potential for altruism is two-fold. First, if the germline and soma are genetically distinct (small *γ*), then cells of the soma place relatively less value on the reproduction of their carrier, and thus are more inclined to promote altruistic behaviours. In contrast, when there is high genetic homogeneity among the germline (high *μ*), then offspring on the patch are more likely to be closely related. This increases relatedness to patchmates and promotes altruism.

Broadly similar patterns occur under other demographic regimes but with some additional complications. For example, if there is additional dispersal before chimeras form, or amongst adults after population regulation, then this further reduces the potential for altruism, reducing the relatedness amongst patchmates but without the concomitant reduction in local competition (see Supplementary Material). Across these cases however, the same essential result still holds, with heterogeneous germlines decreasing the potential for altruism and somatic-germline discordance promoting it.

### Chimerism and intra-individual and intragenomic conflict

Typically, chimerism is thought to foster conflict as cell lineages within individuals compete to increase their own direct representation in the gametes. This situation, wherein different cell lineages jockey for greater numerical representation, is a form of what Patten et al. (2023) labelled “transmission distortion”. However, other forms of internal conflict may also occur within organisms. If different gene copies or cell lineages within an organism have different inclusive fitness interests, then they may benefit from the organism expressing different phenotypes, or “trait distortion” (see also Scott & West, 2019).

In the previous section we treated the soma as a single, well-mixed entity, but in many organisms the soma is highly structured, with different tissues and organs descending from different pools of cells. These distinct developmental histories may result in tissues across the soma not being homogenously related to each other. Instead, some tissues may systematically be more closely, or more distantly, related to the germline. Whereas before we considered a single value of *r*_*s*_, we may need to make room for different *r*_*s*_ for different tissues and organs. For example, in the marmoset, the fused placenta shared by two co-twins is a roughly 50:50 mixture of the two genotypes (Chambers & Hearn, 1985), whereas the brain comprises a mixture of neurons and glia, which are clonal, and cells deriving from the hematopoietic lineage, which can be as mixed as the placenta (Sweeney *et al*., 2012; del Rosario *et al*., 2024).

One consequence of this spatial variation in relatedness is that when we consider social behaviours controlled by different organs, selection may act in different directions upon them. A tissue which is less closely related to the germline (lower *r*_*s*_)—for example, one heavily populated by cells of the hematopoietic lineage in marmosets—will thus have an effectively higher relatedness to social partners (higher 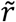), and thus favour more altruistic behaviours. In contrast, a tissue which is more closely related to the germline (higher *r*_*s*_), and thus effectively less related to social partners (lower 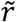), will favour more selfish behaviour (Figure 3). This conflict over control of the phenotype, which arises in the presence of social interactions, is distinct from previously identified internal conflicts of chimeras (e.g. Buss 1987), which take place over numerical representation within the developing body.

**Figure 3:**
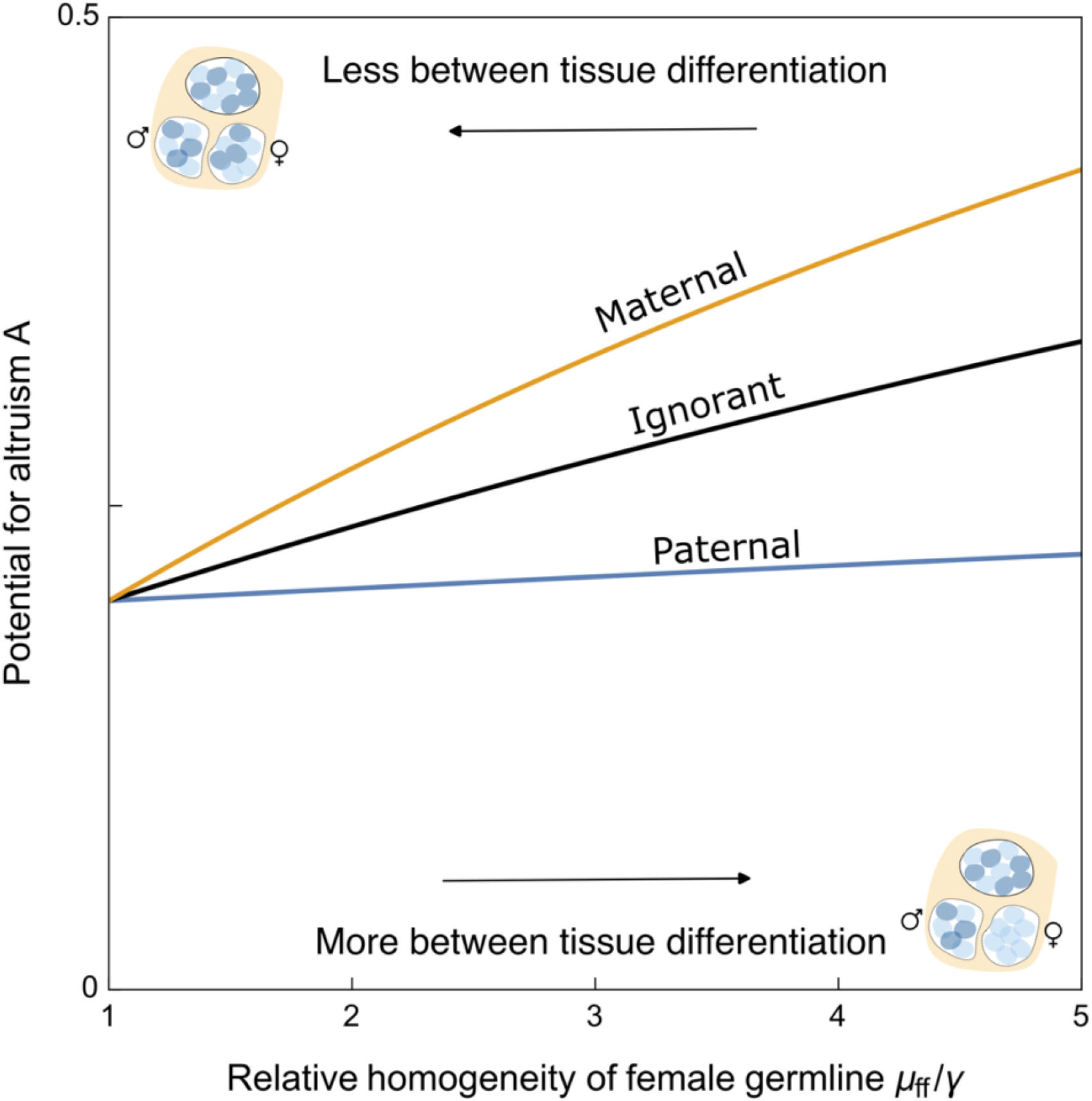
Increasing asymmetry between the homogeneity of male and female germlines generates asymmetries in the potential for altruism between maternal-origin and paternal-origin genes. In the plotted example female germlines are always more homogenous than male germlines, *μ*_mm_=*μ*_fm_=*γ, N*=1/5, *γ*=1/5, and *h*=0.

Alongside fomenting conflicts between cells and organs, chimerism may also be a novel source of conflict between maternal-origin and paternal-origin genes. We now allow for separate male and female germlines within our individuals, which may vary in their degree of genetic homogeneity (*μ*_ff_ ≠ *μ*_mm_). If female germlines are more homogeneous than male germlines (*μ*_ff_>*μ*_mm_) then there are effectively fewer mothers on the patch than fathers.

This results in higher relatedness through maternal-origin genes than through paternal-origin genes. The reverse applies if male germlines are more homogeneous than female germlines (*μ*_ff_ < *μ*_mm_). If we express *μ*_ff_ =*μ*(1 +*δ*) and *μ*_*mm*_=*μ*, assuming that the germlines are no more descended from one another than the soma (*μ*_fm_ =*γ*), then we can compute the relative relatedness to patchmates through maternal-origin and paternal-origin routes as:

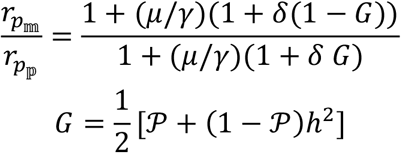

So as *δ*increases—i.e. female germlines are more homogeneous—the numerator increases more than the denominator, and the relatedness through matrilines is relatively higher. (As the maximum value *G*can take is 1/2.)

This asymmetry, in turn, can lead to distinct optima for these two suites of genes, with the gene copies descending from the more homogeneous germline relatively pre-disposed to altruism, whilst those from the more heterogeneous germline relatively disinclined to altruism (Figure 3). Thus, even in the absence of sex differences in ecology—such as dispersal and reproductive skew—or asymmetric transmission genetics—such as X chromosomes or haplodiploidy—sex differences in the degree of germline chimerism are sufficient to generate asymmetries in the relatedness through maternal-origin and paternal-origin genes, and potentially drive the evolution of genomic imprinting (Haig, 2000).

## Discussion

Here, we have developed new theory covering interindividual interactions between chimeric individuals, showing how their unique genetic composition leads to distinct expectations for between-organism patterns of social behaviours. We have shown how the structuring of genetic variation within individuals plays a central role in modulating the potential for altruistic behaviours to evolve, with homogenous germlines and soma-germline differentiation leading to more permissive conditions for altruistic variants to invade. Additionally, such within-organism genetic structure can generate a novel type of internal conflict between cell lineages and tissues over social traits. Those tissues less related to the germline are expected to be relatively altruistic, and those tissues more related to germlines relatively selfish. Finally, in chimeric individuals, differences in the development of male and female germlines can lead to systematic differences in maternal and paternal relatedness coefficients, an asymmetry that may drive the evolution of genomic imprinting.

One key factor our model has highlighted is the importance of genetic structure within chimeric individuals—specifically, the relationships between the cells within the germline (*μ*) and between the germline and different somatic tissues (*γ*). In some species, experimental work has begun to identify patterns in how initial genetic heterogeneity is distributed through development. For example: in macroalgae, bicolour chimeras show that the multilineage holdfasts typically develop separate, uniclonal branches (Santelices *et al*., 2017); in sponges, by using different fluorescent dyes for separate cell lineages, it was found that they form distinct tissue types (Gauthier & Degnan, 2008); whilst in corals spatial genotyping has demonstrated that branches maintain cellular diversity through the organism (Guerrini *et al*., 2021). Despite this, improved resolution experiments—across more species and in more natural settings—are needed to draw generalisations about patterns of typical internal relatedness patterns and their developmental and ecological determinants in these groups. With the increasing affordability of single cell sequencing and improvements in cell lineage tracking, this may soon become a realistic goal for many of these non-model organisms (Wagner & Klein, 2020; Alfieri *et al*., 2022).

One recent, exciting example of such approaches was shown in marmosets, which have long been known to have chimerism due to cell sharing between siblings with fused placentas (Hill, 1932; Wislocki, 1939; Benirschke *et al*., 1962; Gengozian *et al*., 1964; Ross *et al*., 2007; Sweeney *et al*., 2012). Using single-nucleus RNA sequencing, del Rosario and colleagues (2024) could infer, for individual cells, both the cell type and the sibling zygote it descended from. With these data, they showed that all observed chimerism arises from the hematopoietic stem cell lineage, with the extent of chimerism varying across different types of descendant cell types. Kuppfer cells (∼15%) (liver specific macrophages) and lymphocytes (4%) had lower degrees of chimerism than B and CD8+ T-cells (∼29% and 32%, respectively). Moreover, microglia, another hematopoietic-derived cell type, showed variable degrees of chimerism dependent on the area of the brain sampled, with much higher fractions of sibling cells in the striatum (56%) as compared to the thalamus (11%). In contrast, neurons and hepatocytes, which do not descend from hematopoietic stem cells, showed no evidence of chimerism. Thus, here there is potential for a variety of intra-tissue conflicts over social traits, with closely interacting cell types showing systematically different relative valuations of self and social partners.

Often the focus for internal conflicts in chimeras has been about transmission, or entry into the germline (Buss, 1987). Here, we have identified a more subtle kind of conflict, where different tissues may disagree about the control of social traits (see also Patten 2021). Whilst early partitioning of the germline might restrict direct conflict over transmission, conflicts over social traits might still emerge if different tissues have different relatedness coefficients to self and social partners. In this way, it mirrors another form of internal conflict, that between maternal-origin and paternal-origin genes (Haig, 2002). Here the conflict is not a matter of jockeying for transmission, for imprinted genes obey Mendel’s laws. Instead in these cases, the two alleles of a locus may come into conflict over the treatment of relatives because of how such behaviors influence their respective indirect fitnesses. Only by considering the full, inclusive fitness of the organism—and, just as importantly, its constituent parts—can these fissures in the adaptive integrity of the organism be observed (Gardner & U beda, 2017; Gardner, n.d.). Indeed, chimerism itself may be a novel source of maternal-origin and paternal-origin relatedness asymmetries— and thus intragenomic conflict—if there are differences in how male and female germlines form.

We have focused an abstract model to capture some of the key social evolutionary consequences of chimerism. Yet, it is important to bear in mind the diversity of forms chimerism may take (Rinkevich, 2011). Chimeric species vary widely in the ecologies they inhabit, their social interactions, and in the kinds and timing of chimerism in relation to other life cycle events. In particular, we have focused on within-generation (“horizontal”) chimerism, in contrast to previous work which has most often focused on intergenerational (“vertical”) chimerism (Haig, 2014b, 2014a; Boddy *et al*., 2015; U beda & Wild, 2023). The challenge for future theoretical work in this area lies in simultaneously developing specific models tailored to the biology of different groups and forms of chimerism, whilst also finding common principles that apply across this diversity.

Finally, whilst we have focused our attention on multicellular organisms, and chimeric mixtures of cell lineages, chimerism-like phenomena can appear at multiple biological scales. Above the individual, social groups composed of genetically heterogeneous individuals may engage in both cooperative and competitive interactions with other social groups (Rodrigues *et al*., 2022, 2023). Pleometrosis, the formation of a joint colony between multiple queens, has been observed in many social insects, including mites, ants, wasps, thrips, aphids, and termites (Bono & Crespi, 2006). This too can be viewed as a higher-level form of chimerism, instead occurring at the level of the colony (or superorganism). At lower levels, the genomes of sexual organisms can be seen as chimeras of maternal and paternal-origin genomes. Paternal genome elimination—a genetic system whereby males receive, but do not transmit, a paternal-origin genome—shares many similarities to the example analysed here. The transmission of only a maternal genome in males makes their gametes more homogenous, but the presence of the paternal genome generates genetic differences between soma and germline. Subsequently, conditions for altruism to evolve are more permissive (Hitchcock & Gardner, 2022).

The major transitions framework views the evolution of individuality as a social-evolutionary problem: how to ensure cooperation and to minimize conflict among the parts of collective entities (Maynard Smith & Szathmáry, 2010; Bourke, 2011; West *et al*., 2015)—in other words, how to overcome Dawkins’ (1990) paradox of the organism (“…that it is not torn apart by its conflicting replicators, but stays together and works as a purposeful entity, apparently on behalf of all of them”). One way to achieve such unity of purpose is to avoid genetic variation within collectives, but this is a strategy that chimeric organisms completely spurn. How chimeric organisms cope with the risk from their internal genetic variation and the various conflicts it foments, some of which are chronicled here for the first time, should spur further theoretical and empirical work.

## Author contributions

TJH first conceived of the research, wrote the first draft, and conducted the analyses. MMP assisted in the conception of the research and helped write the manuscript.

## Data and code availability statement

There is no associated data. Full methods to recreate results can be found in the main text and supporting Supplementary Material.

## Acknowledgments and Funding

This work is supported by the Special Postdoctoral Researchers Program funded by RIKEN (TJH), and the John Templeton Foundation (MMP, grant #63320).

