## Supplementary Material for "Chimerism and altruism"

#### 1 Life-cycle and consanguinities

##### 1.1 Life cycle

We consider a population hierarchically structured in the following way. There are an infinite number of patches, on those patches there are initially $N$ adults, who are composed of potentially multiple cell lineages. We assume that parents on each patch mate and produce a large number of zygotes, with each patch having the same same intrinsic fecundity. Those zygotes may then disperse - dispersal phase 1 - and a fraction remain on the patch with probability $h_1$. Zygotes then aggregate to form individual organisms. We assume that the aggregations are not too large, such that the number of chimeric organisms that now exist on the patch $n$ is still large enough that we can ignore terms of $(1/n)^2$. There is then a phase of growth of those individuals, distributing cell lineages across the body. Individuals may then disperse - dispersal phase 2 - with a fraction $h_2$ remaining on the patch. There is then density dependent regulation, with $N$ adult breeders remaining on the patch. There is then another round of dispersal - dispersal phase 3 - where individuals remain on their natal patch with probability $h_3$. Those individuals then mate, and the life-cycle begins again.

##### 1.2 Consanguinities

We track consanguinities amongst the set of zygotes produced on a patch at the start of the life cycle. We follow three variables: the consanguinity between maternal-origin genes $Q_{\text{mm}}$, the consanguinity between paternal-origin genes $Q_{\text{pp}}$, and the consanguinity between maternal-origin and paternal-origin genes $Q_{\text{mp}}$. We can write these out in vector form:

$$\mathbf{Q} = \begin{pmatrix} Q_{\text{mm}} \\ Q_{\text{mp}} \\ Q_{\text{pp}} \end{pmatrix} \tag{S1}$$

The consanguinities in the next generation are $\mathbf{Q}'$. We describe the relation between them as follows.

$$Q'_{\text{mm}} = \mathscr{P}_{\text{ff}}\mu_{\text{ff}}Q_{\text{c}} + \left(\mathscr{P}_{\text{ff}}(1-\mu_{\text{ff}})h_1^2 + (1-\mathscr{P}_{\text{ff}})h_1^2h_2^2h_3^2\right)Q_{\text{p}} \tag{S2a}$$

$$Q'_{\mathrm{mp}} = \mathscr{P}_{\mathrm{fm}}\mu_{\mathrm{fm}}Q_{\mathrm{c}} + \left(\mathscr{P}_{\mathrm{fm}}(1-\mu_{\mathrm{fm}})h_1^2 + (1-\mathscr{P}_{\mathrm{fm}})h_1^2h_2^2h_3^2\right)Q_{\mathrm{p}} \tag{S2b}$$

$$Q'_{\mathrm{pp}} = \mathscr{P}_{\mathrm{mm}}\mu_{\mathrm{mm}}Q_{\mathrm{c}} + \left(\mathscr{P}_{\mathrm{mm}}(1-\mu_{\mathrm{mm}})h_1^2 + (1-\mathscr{P}_{\mathrm{mm}})h_1^2h_2^2h_3^2\right)Q_{\mathrm{p}} \tag{S2c}$$

Where:

$$Q_{\mathrm{c}} = \alpha^2 + 2\alpha(1-\alpha)Q_{\mathrm{mp}} + (1-\alpha)^2 \tag{S3a}$$

$$Q_{\mathrm{p}} = \alpha^2 Q_{\mathrm{mm}} + 2\alpha(1-\alpha)Q_{\mathrm{mp}} + (1-\alpha)^2 Q_{\mathrm{pp}} \tag{S3b}$$

Where $\alpha$ is the probability of sampling a maternal-origin gene copy, and hence $1-\alpha$ the probability of sampling a paternal-origin gene copy, $\mu_{\mathrm{ij}}$ is the probability that two cells sampled from the gonads of sex $i$ and $j$ from a single individual are from the same cell lineage, and $\mathscr{P}_{\mathrm{ij}}$ is the probability of sibship through sex routes $i$ and $j$.

We can collate this information into the matrix **S** describing the transition probabilities between consanguinities each generation, and the vector $\mathbf{S}_c$ which describes the increase in consanguinity due to coalescence. We can then solve for when $\mathbf{Q}' = \mathbf{Q}$:

$$\mathbf{Q} = \mathbf{S}.\mathbf{Q} + \mathbf{S}_c \tag{S4}$$

Full expressions are unwieldy, however, if we assume no differences in reproduction $\mathscr{P}_{\mathrm{ij}} = \mathscr{P}$, that there is a single germline $\mu_{\mathrm{ij}} = \mu$, and that dispersal only happens amongst juvenile chimeras ($h_1 = h_3 = 0$, $h_2 = h$) then:

$$\mathbf{Q}_{\mathrm{mm}} = \mathbf{Q}_{\mathrm{mp}} = \mathbf{Q}_{\mathrm{pp}} = \frac{\mathscr{P}\mu}{\mathscr{P}\mu + 2(1-\mathscr{P})\left(1-h^2\right)} \tag{S5}$$

This is the scenario we focus on in the main text. We can also then rewrite the consanguinities between gene copies within the cell ($Q_{\mathrm{c}}$), or between other zygotes on the patch ($Q_{\mathrm{p}}$) in terms of our demographic parameters.

##### 1.3 Consanguinities between germline and soma

The consanguinity between the germline and soma can be described in the following way. If we assume that there is some asexual, developmental process that occurs as the organism develops - similar to other forms of asexual proliferation - then we can describe the probability that a gene copy in the germline and in the soma come from the same initial cell lineage as $\gamma_{\mathrm{i}}$, where $i$ can stand for either sex (Prugnolle et al., 2005). In this case this means that the consanguinity between the soma and the female germline is:

$$Q^{o}_{\mathrm{s,g_f}} = \gamma_{\mathrm{f}}Q_{\mathrm{c}} + (1-\gamma_{\mathrm{f}})h_1^2 Q_{\mathrm{p}} \tag{S6a}$$

And between the soma and the male germline:

$$Q^{o}_{\mathrm{s,g_m}} = \gamma_{\mathrm{m}}Q_{\mathrm{c}} + (1-\gamma_{\mathrm{m}})h_1^2 Q_{\mathrm{p}} \tag{S6b}$$

We can then further partition this to consider consanguinities between specific pairs of genes. For example, if we consider the consanguinities between genes of different parental-origin (either maternal-origin $\mathbb{m}$ or paternal-origin $\mathbb{p}$) to the cell or to the patch then:

$$Q_{\mathrm{c}_{\mathbb{m}}} = \alpha + (1-\alpha)Q_{\mathrm{mp}} \tag{S7a}$$

$$Q_{\mathrm{p}_{\mathbb{m}}} = \alpha Q_{\mathrm{mm}} + (1-\alpha)Q_{\mathrm{mp}} \tag{S7b}$$

$$Q_{\mathrm{c}_{\mathbb{p}}} = \alpha Q_{\mathrm{mp}} + (1-\alpha) \tag{S7c}$$

$$Q_{\mathrm{p}_{\mathbb{p}}} = \alpha Q_{\mathrm{mp}} + (1-\alpha)Q_{\mathrm{pp}} \tag{S7d}$$

Thus, this gives us the following consanguinities:

$$Q^{o}_{\mathrm{s}_{\mathbb{m}},\mathrm{g_f}} = \gamma_{\mathrm{f}} Q_{\mathrm{c}_{\mathbb{m}}} + (1-\gamma_{\mathrm{f}})h_1^2 Q_{\mathrm{p}_{\mathbb{m}}} \tag{S8a}$$

$$Q^{o}_{\mathrm{s}_{\mathbb{p}},\mathrm{g_f}} = \gamma_{\mathrm{f}} Q_{\mathrm{c}_{\mathbb{p}}} + (1-\gamma_{\mathrm{f}})h_1^2 Q_{\mathrm{p}_{\mathbb{p}}} \tag{S8b}$$

$$Q^{o}_{\mathrm{s}_{\mathbb{m}},\mathrm{g_m}} = \gamma_{\mathrm{m}} Q_{\mathrm{c}_{\mathbb{m}}} + (1-\gamma_{\mathrm{m}})h_1^2 Q_{\mathrm{p}_{\mathbb{m}}} \tag{S8c}$$

$$Q^{o}_{\mathrm{s}_{\mathbb{p}},\mathrm{g_m}} = \gamma_{\mathrm{m}} Q_{\mathrm{c}_{\mathbb{p}}} + (1-\gamma_{\mathrm{m}})h_1^2 Q_{\mathrm{p}_{\mathbb{p}}} \tag{S8d}$$

#### 2 Inclusive fitness analysis

First, we perform an inclusive fitness analysis (Wild and Scott, 2023). We consider two different types of actor, maternal-origin genes in somatic tissue ($s_{\mathbb{m}}$) and paternal-origin genes in somatic tissue ($s_{\mathbb{p}}$). Thus the condition for our trait to increase can be written as:

$$\Delta W_{\mathrm{IF}} = \Delta W_{\mathrm{IF}}^{s_{\mathbb{m}}} + \Delta W_{\mathrm{IF}}^{s_{\mathbb{p}}} > 0 \tag{S9}$$

We assign a fraction $\rho$ of the control over the phenotype to maternal-origin genes, and a fraction $1-\rho$ to paternal origin genes. The deviant strategy imposes a fitness cost of $C$ on the focal individual and provides a benefit $B$ on a random member of the focal patch.

The reproductive value of an individual through the sex $i$ germline is $v_{g_i}$, where $i$ could be either female $f$ or male $m$. The relatedness between our somatic actors, and germline recipients is $r^{k}_{s_{\mathbb{j}}\to \mathrm{g_i}}$, where $i$ is the sex of the germline (either female or male), $j$ is the parent-of-origin of the genetic actor, and is either within the same focal organism $o$, or to a random individual on the patch $p$.

The condition for increase is:

$$\begin{aligned} &\rho\left[-C\left(v_{\mathrm{g_f}} r^{o}_{s_{\mathbb{m}}\to \mathrm{g_f}} + v_{\mathrm{g_m}} r^{o}_{s_{\mathbb{m}}\to \mathrm{g_m}}\right) + B\left(v_{\mathrm{g_f}} r^{p}_{s_{\mathbb{m}}\to \mathrm{g_f}} + v_{\mathrm{g_m}} r^{p}_{s_{\mathbb{m}}\to \mathrm{g_m}}\right)\right] \\ &+ (1-\rho)\left[-C\left(v_{\mathrm{g_f}} r^{i}_{s_{\mathbb{p}}\to \mathrm{g_f}} + v_{\mathrm{g_m}} r^{o}_{s_{\mathbb{p}}\to \mathrm{g_m}}\right) + B\left(v_{\mathrm{g_f}} r^{p}_{s_{\mathbb{p}}\to \mathrm{g_f}} + v_{\mathrm{g_m}} r^{p}_{s_{\mathbb{p}}\to \mathrm{g_m}}\right)\right] > 0 \end{aligned} \tag{S10}$$

Writing the costs and benefits in terms of marginal survival costs $-c$ and marginal survival benefits $b$, and accounting for the scale of local competition $a$, will be:

$$C = c \tag{S11a}$$

$$B = b - a(b-c) \tag{S11b}$$

Noting that the probability the maternal-origin gene is transmitted is $\alpha$, then the reproductive values for gene copies in female germlines will be $v_{\mathrm{g_f}} = c_{\mathrm{g_f}}/n = \alpha/n$, and thus for gene copies in male germlines $v_{\mathrm{g_m}} = c_{\mathrm{g_m}}/n =$

$(1-\alpha)/n$. Using this, and as the effect on the two germlines is equivalent, then we can treat the germline as a single recipient of the behaviour, where we have averaged over their reproductive values. In this case the relatedness coefficients will be:

$$r^o_{s_{\mathbb{m}}\to g} = \left(\alpha r^o_{s_{\mathbb{m}}\to g_{\mathrm{f}}} + (1-\alpha) r^o_{s_{\mathbb{m}}\to g_{\mathrm{m}}}\right) = (\alpha\gamma_{\mathrm{f}} + (1-\alpha)\gamma_{\mathrm{m}})Q_{\mathrm{c}_{\mathbb{m}}} + h_1^2(\alpha(1-\gamma_{\mathrm{f}}) + (1-\alpha)(1-\gamma_{\mathrm{m}}))Q_{\mathrm{p}_{\mathbb{m}}} \tag{S12a}$$

$$r^p_{s_{\mathbb{m}}\to g} = \left(\alpha r^p_{s_{\mathbb{m}}\to g_{\mathrm{f}}} + (1-\alpha) r^p_{s_{\mathbb{m}}\to g_{\mathrm{m}}}\right) = h_1^2 Q_{\mathrm{p}_{\mathbb{m}}} \tag{S12b}$$

$$r^o_{s_{\mathbb{p}}\to g} = \left(\alpha r^o_{s_{\mathbb{p}}\to g_{\mathrm{f}}} + (1-\alpha) r^o_{s_{\mathbb{p}}\to g_{\mathrm{p}}}\right) = (\alpha\gamma_{\mathrm{f}} + (1-\alpha)\gamma_{\mathrm{m}})Q_{\mathrm{c}_{\mathbb{p}}} + h_1^2(\alpha(1-\gamma_{\mathrm{f}}) + (1-\alpha)(1-\gamma_{\mathrm{m}}))Q_{\mathrm{p}_{\mathbb{p}}} \tag{S12c}$$

$$r^p_{s_{\mathbb{p}}\to g} = \left(\alpha r^p_{s_{\mathbb{p}}\to g_{\mathrm{f}}} + (1-\alpha) r^p_{s_{\mathbb{p}}\to g_{\mathrm{m}}}\right) = h_1^2 Q_{\mathrm{p}_{\mathbb{p}}} \tag{S12d}$$

And putting this all together we can rewrite out condition for increase as:

$$\frac{c}{b} < \frac{(1-a)\left(\rho r^p_{s_{\mathbb{m}}\to g} + (1-\rho) r^p_{s_{\mathbb{p}}\to g}\right)}{\left(\rho r^o_{s_{\mathbb{m}}\to g} + (1-\rho) r^o_{s_{\mathbb{p}}\to g}\right) - a\left(\rho r^p_{s_{\mathbb{m}}\to g} + (1-\rho) r^p_{s_{\mathbb{p}}\to g}\right)} = \frac{(1-a)\, r_{\mathrm{p}}}{r_{\mathrm{o}} - a r_{\mathrm{p}}} \tag{S13}$$

Where the $r_{\mathrm{o}}$ and $r_{\mathrm{p}}$ are the appropriately averaged relatedness coefficients incorporating both the asymmetric transmission (and thus reproductive value) of some gene copies, and the asymmetric expression (and thus phenotypic control) of others.

### 3 Neighbour-modulated fitness approach

We now perform a direct fitness, or neighbour-modulated fitness approach (Taylor, 1996; Taylor and Frank, 1996; Frank, 1998; Rousset, 2004). We will census the life-cycle after aggregation. We first write out the fitness functions describing the number of offspring a focal individual produces after one iteration of the life cycle. The probability of survival of our focal chimeric individual $S(x, y)$, which is a function of their own somatic trait value $x$ and the trait value of their social partners $y$. Individuals then either disperse, competing with unrelated individuals, or remain on their focal patch, competing with a proportion of related neighbours. They then produce a large number of offspring $\kappa$.

$$w(x,y,z) = h_2\left(\frac{S(x,y)}{h_2 S(y,y) + (1-h_2)S(z,z)}\right)\kappa + (1-h_2)\left(\frac{S(x,y)}{S(z,z)}\right)\kappa \tag{S14a}$$

$$W(x,y,z) = \frac{w(x,y,z)}{w(z,z,z)} \tag{S14b}$$

We can then write:

$$\frac{1}{S(z,z)}\frac{\partial S(x,y)}{\partial x}\bigg|_{x=y=z} = -c \tag{S15a}$$

$$\frac{1}{S(z,z)}\frac{\partial S(x,y)}{\partial y}\bigg|_{x=y=z} = b \tag{S15b}$$

$$\frac{\partial W}{\partial x}\bigg|_{x=y=z} = -c \tag{S15c}$$

$$\frac{\partial W}{\partial y}\bigg|_{x=y=z} = b - h_2^2(b-c) = b - a(b-c) \tag{S15d}$$

We then write out the condition for our trait to increase. We first partition fitness into male and female components, weighting by their respective class reproductive values $V_{\mathrm{f}}$ and $V_{\mathrm{m}}$. We then partition the change in fitness due to ones own phenotype, and due to changes in the phenotype of social partners, weighting by the appropriate relatedness coefficients.

$$
\begin{aligned}
\frac{dW}{dg} &= V_{\mathrm{f}}\frac{dW_{\mathrm{f}}}{dg_{\mathrm{f}}} + V_{\mathrm{m}}\frac{dW_{\mathrm{m}}}{dg_{\mathrm{m}}} \\
&= V_{\mathrm{f}}\left(\frac{\partial W_{\mathrm{f}}}{\partial x}\frac{dx}{dg_{\mathrm{f}}} + \frac{\partial W_{\mathrm{f}}}{\partial y}\frac{dy}{dg_{\mathrm{f}}}\right) + V_{\mathrm{m}}\left(\frac{\partial W_{\mathrm{m}}}{\partial x}\frac{dx}{dg_{\mathrm{m}}} + \frac{\partial W_{\mathrm{m}}}{\partial y}\frac{dy}{dg_{\mathrm{m}}}\right) \\
&= V_{\mathrm{f}}\left(\frac{\partial W_{\mathrm{f}}}{\partial x}r^{o}_{g_{\mathrm{f}}\to s} + \frac{\partial W_{\mathrm{f}}}{\partial y}r^{p}_{g_{\mathrm{f}}\to s}\right) + V_{\mathrm{m}}\left(\frac{\partial W_{\mathrm{m}}}{\partial x}r^{o}_{g_{\mathrm{m}}\to s} + \frac{\partial W_{\mathrm{m}}}{\partial y}r^{p}_{g_{\mathrm{m}}\to s}\right) \\
&= V_{\mathrm{f}}\left(-cr^{o}_{g_{\mathrm{f}}\to s} + (b-a(b-c))r^{p}_{g_{\mathrm{f}}\to s}\right) + V_{\mathrm{m}}\left(-cr^{o}_{g_{\mathrm{m}}\to s} + (b-a(b-c))r^{p}_{g_{\mathrm{m}}\to s}\right) \\
&> 0
\end{aligned}
\tag{S16}
$$

We can further split the relatedness coefficients by which gene in the soma has control over the phenotype:

$$r^{o}_{g_{\mathrm{f}}\to s} = \rho r^{o}_{g_{\mathrm{f}}\to s_{\mathbb{m}}} + (1-\rho)r^{o}_{g_{\mathrm{f}}\to s_{\mathbb{p}}} \tag{S17a}$$

$$r^{o}_{g_{\mathrm{m}}\to s} = \rho r^{o}_{g_{\mathrm{m}}\to s_{\mathbb{m}}} + (1-\rho)r^{o}_{g_{\mathrm{m}}\to s_{\mathbb{p}}} \tag{S17b}$$

$$r^{p}_{g_{\mathrm{f}}\to s} = \rho r^{p}_{g_{\mathrm{f}}\to s_{\mathbb{m}}} + (1-\rho)r^{p}_{g_{\mathrm{f}}\to s_{\mathbb{p}}} \tag{S17c}$$

$$r^{p}_{g_{\mathrm{m}}\to s} = \rho r^{p}_{g_{\mathrm{m}}\to s_{\mathbb{m}}} + (1-\rho)r^{p}_{g_{\mathrm{m}}\to s_{\mathbb{p}}} \tag{S17d}$$

If we make the substitution that $V_{\mathrm{f}} = \alpha$ and $V_{\mathrm{m}} = 1-\alpha$, and then rearrange, we get the following expression for increase:

$$\frac{c}{b} < \frac{(1-a)\left(\alpha r^{o}_{g_{\mathrm{f}}\to s} + (1-\alpha)r^{o}_{g_{\mathrm{m}}\to s}\right)}{\left(\alpha r^{o}_{g_{\mathrm{f}}\to s} + (1-\alpha)r^{o}_{g_{\mathrm{m}}\to s}\right) + a\left(\alpha r^{o}_{g_{\mathrm{f}}\to s} + (1-\alpha)r^{o}_{g_{\mathrm{m}}\to s}\right)} = \frac{(1-a)\,r_{\mathrm{p}}}{r_{\mathrm{o}} - ar_{\mathrm{p}}} \tag{S18}$$

We can see that this generates the same result as in our inclusive fitness analysis as:

$$
\begin{aligned}
r_{o} &= \alpha\left(r^{o}_{g_{\mathrm{f}}\to s}\right) + (1-\alpha)\left(r^{o}_{g_{\mathrm{f}}\to s}\right) \\
&= \alpha\left(\rho r^{o}_{g_{\mathrm{f}}\to s_{\mathbb{m}}} + (1-\rho)r^{o}_{g_{\mathrm{f}}\to s_{\mathbb{p}}}\right) + (1-\alpha)\left(\rho r^{o}_{g_{\mathrm{m}}\to s_{\mathbb{m}}} + (1-\rho)r^{o}_{g_{\mathrm{m}}\to s_{\mathbb{p}}}\right) \\
&= \rho\left(\alpha r^{o}_{g_{\mathrm{f}}\to s_{\mathbb{m}}} + (1-\alpha)r^{o}_{g_{\mathrm{m}}\to s_{\mathbb{m}}}\right) + (1-\rho)\left(\alpha r^{o}_{g_{\mathrm{f}}\to s_{\mathbb{p}}} + (1-\alpha)r^{o}_{g_{\mathrm{m}}\to s_{\mathbb{p}}}\right) \\
&= \rho\left(\alpha r^{o}_{s_{\mathbb{m}}\to g_{\mathrm{f}}} + (1-\alpha)r^{o}_{s_{\mathbb{m}}\to g_{\mathrm{m}}}\right) + (1-\rho)\left(\alpha r^{o}_{s_{\mathbb{p}}\to g_{\mathrm{f}}} + (1-\alpha)r^{o}_{s_{\mathbb{p}}\to g_{\mathrm{m}}}\right) \\
&= \rho\left(r^{o}_{s_{\mathbb{m}}\to g}\right) + (1-\rho)\left(r^{o}_{s_{\mathbb{p}}\to g}\right)
\end{aligned}
\tag{S19}
$$

Writing $\tilde{r} = r_{\mathrm{p}}/r_{\mathrm{o}}$, then:

$$\frac{c}{b} < \frac{(1-a)\tilde{r}}{1-\tilde{r}} \tag{S20}$$

### 4 Additional phases of dispersal

In the main text we focus on the scenario where there is one phase of dispersal after the social interaction, but before density regulation. We consider the effects of different timings of dispersal relative to these other effects here. If we ignore any sex-specific differences in reproductive skew or germline development, but allow for various timings of dispersal then the potential for altruism is:

$$A = \frac{h_1^2\left(1-h_2^2\right)\mathscr{P}\mu}{\gamma\left(1-h_1^2\left(\mathscr{P} - h_2^2h_3^2(1-\mathscr{P})\right)\right) + h_1^2\left(1-h_2^2\right)\mathscr{P}\mu} \tag{S21}$$
